# Adaptive Spike-Artifact Removal from Local Field Potentials Uncovers Prominent Beta and Gamma Band Neuronal Synchronization

**DOI:** 10.1101/820571

**Authors:** Kianoush Banaie Boroujeni, Paul Tiesinga, Thilo Womelsdorf

## Abstract

**Background:** Many neurons synchronize their action potentials to the phase of local field potential (LFP) fluctuations in one or more frequency bands. Analyzing this spike-to-LFP synchronization is challenging, however, when neural spikes and LFP are generated in the same local circuit, because the spike’s action potential waveform leak into the LFP and distort phase synchrony estimates. Existing approaches to address this spike bleed-through artifact relied on removing the average action potential waveforms of neurons, but this leaves artifacts in the LFP and distorts synchrony estimates.

**New Method:** We describe a spike-removal method that surpasses these limitations by decomposing individual action potentials into their frequency components before their removal from the LFP. The adaptively estimated frequency components allow for variable spread, strength and temporal variation of the spike artifact.

**Results:** This adaptive approach effectively removes spike bleed-through artifacts in synthetic data with known ground truth, and in single neuron and LFP recordings in nonhuman primate striatum. For a large population of neurons with both narrow and broad action potential waveforms, the use of adaptive artifact removal uncovered 20-35 Hz beta and 35-45 Hz gamma band spike-LFP synchronization that would have remained contaminated otherwise.

**Comparison with Existing Methods:** We demonstrate that adaptive spike-artifact removal cleans LFP data that remained contaminated when applying existing Bayesian and non-Bayesian methods of average spike-artifact removal.

**Conclusions:** Applying adaptive spike-removal from field potentials allows to estimate the phase at which neurons synchronize and the consistency of their phase-locked firing for both beta and low gamma frequencies. These metrics may prove essential to understand cell-to-circuit neuronal interactions in multiple brain systems.

## 1. Introduction

Spiking activity of many different neuron types synchronize to the local field potential (LFP) (Klausberger and Somogyi, 2008; Roux and Buzsaki, 2015). The strength of this spike-LFP phase synchronization can predict how a neuron contributes to the functioning of the circuit (Pesaran et al., 2018; Womelsdorf and Everling, 2015), how strong it gates afferent inputs through the circuit (Cardin et al., 2009; Tiesinga et al., 2008; Womelsdorf et al., 2014b), and how strong it communicates with upstream brain areas (Bastos et al., 2015; Fries, 2015; Womelsdorf et al., 2007). The importance of spike-LFP relationships is highlighted by reports that the likelihood of spiking responses can sometimes be better predicted by the phase of the local field than by spiking of neurons at nearby electrodes (Besserve et al., 2010). One reason for the informativeness of the LFP is that it is generated by currents flowing along dendritic and axonal membranes (Einevoll et al., 2013; Mitzdorf, 1985; Reimann et al., 2013; Schomburg et al., 2012). When these transmembrane currents reflect synaptic inputs or subthreshold depolarization levels they can effectively set a gain control for the spike output of neurons and thereby provide crucial insights into information processing (Azouz and Gray, 2003; Fries, 2005; Womelsdorf et al., 2014b).

However, the close relationship of spiking activity to the LFP can be artifactual when the local field potential activity is contaminated by the action potentials of the spiking neurons themselves as opposed to be based on transmembrane currents from the local circuit (Buzsaki et al., 2012; Einevoll et al., 2013; Ray, 2015). Such contamination of the LFP can occur for spiking activity of neurons in the immediate surroundings (within ∼200 μm) of a recorded neuron (Watson et al., 2018) and becomes evident as spike-bleed through artifacts in spike-triggered LFP averages or as artifactual spike-LFP synchrony, because the LFP phase of the spiking activity is contaminated by the spike itself and thus is not informative beyond the spike time itself. This spike bleed through effect is not only evident for the high frequencies at which the fast action potentials main power reside (around ∼0.8-5 kHz), but it can dominate spike-LFP measurements down to ∼100 Hz, with lower but discernible contributions for frequencies as low as 25Hz (Ardid et al., 2015; Ray, 2015; Schomburg et al., 2012; Zanos et al., 2011).

To prevent the spike bleed through effects, a clean LFP has to be estimated. In many studies this is achieved by measuring the LFP from electrode tips that are far (>200 μm) away from the electrode recording the spiking neuron so that it does not affect the potential. This approach might be successful when the LFP is homogeneous over several hundreds of micrometers, However, there are major limitations to this approach in many settings. Firstly, the LFP might not be homogeneous between two electrodes that are >200 μm apart, but rather changes its phase systematically as quantified e.g. in traveling waves. Secondly, it is unclear how much distance between spike- and LFP electrode should be considered safe. For example, with silicon shank electrodes in somatosensory cortex there are apparent spike-bleed through artifacts evident at ∼≥100 Hz spike-LFP wavelet spectra with channels 200 μm away (e.g. see Figure 6, Suppl. 3 and 11 in (Watson et al., 2018)). This is a problem when considering that local cell-to-circuit interactions between neurons can be limited to a <150 μm diameter with spike triggered averages being essentially flat at larger distances of the neuron to the site of LFP recording (Fujisawa et al., 2008).

These considerations suggest that a versatile approach to estimate the LFP around the spiking neuron is to clean the LFP activity from influences of the spike of a neuron itself. Existing approaches for removing spike artifacts have either focused on the average waveform of a neuron to subtract its average contribution over an arbitrarily defined time window around the spike time (Pesaran et al., 2002; Zanos et al., 2012; Zanos et al., 2011), estimated the spike through a dictionary based marching pursuit algorithm (Ray et al., 2008), or designed an on-average optimal linear filter predicting spiking influences on the LFP (David et al., 2010), or removed data and interpolated across the time around the spike occurrence to remove the spikes influence on spectral estimates of spike-LFP synchrony (Ardid et al., 2015; Galindo-Leon and Liu, 2010; Okun et al., 2010; Womelsdorf et al., 2010). However, all of these approaches have in common that they do leave residual spike-artifacts in the data in those lower 25-100 Hz frequencies that contain physiologically interesting information (Pesaran et al., 2018).

Here, we report of a novel approach that removes spike-related transients in low frequencies of the LFP by addressing several limitations of prior approaches. First, in contrast to existing methods the novel approach does not require an *a priori* determination of a short duration that should be taken into account to include in the artifact estimation. This is important to be able to remove slower spike accompanying events (e.g. slow after-hyperpolarization, plateau potentials or preceding EPSC barrages of activated nearby synapses) as sources of the spike-artifact. Secondly, the proposed method estimates the peak time and duration for individual spikes and is not based on a grand average spike waveform and thus allows for variable width and height (i.e. shape) of action potential waveforms. This feature is critically important to allow for changes in spike amplitude and shape that occur e.g. for burst firing neurons. By incorporating these features, we show in ground truth simulations that our approach effectively removes the action potential contributions to the LFP without introducing distortions of the phase that prior methods could not avoid. We then show in electrophysiological recordings in nonhuman primate striatum that removal of spike-transients is essential to detect narrow band spike-LFP phase synchrony in the beta (20-30 Hz) and gamma (35-45 Hz) bands. Notably, phase synchrony was reliably estimated for neurons with narrow as well as broad waveforms, suggesting that the adaptive spike-artifact removal approach allows distinguishing how different cell classes synchronize to the local LFP.

## 2. Materials and Methods

### 2.1 Electrophysiological Recording

Data was collected from two male rhesus macaques (Macaca mulatta) from the head of the caudate and the ventral striatum as described in full in Oemisch et al, (2019). All animal care and experimental protocols were approved by the York University Council on Animal Care and were in accordance with the Canadian Council on Animal Care guidelines. Extra-cellular recordings were made with tungsten electrodes (impedance 1.2 - 2.2 MOhm, FHC, Bowdoinham, ME) through rectangular recording chambers implanted over the right hemisphere. Electrodes were lowered daily through guide tubes using software-controlled precision micro-drives (NAN Instruments Ltd., Israel). Wideband local field potential (LFP) data was recorded with a multi-channel acquisition system (Digital Lynx SX, Neuralynx) with a 32kHz sampling rate. Spiking activity was obtained following a 300 - 8000 Hz passband filter and further amplification and digitization at a 32 kHz sampling rate. Sorting and isolation of single unit activity was performed offline with Plexon Offline Sorter, based on the first two principal components of the spike waveforms and the temporal stability of isolated neurons. Only well isolated neurons were considered for analysis (Ardid et al., 2015). All wideband local field analysis was done with the 32kHz sampled data if not otherwise explicitly stated. Experiments were performed in a custom-made sound attenuating isolation chamber. Monkeys sat in a custom-made primate chair viewing visual stimuli on a computer monitor (60Hz refresh rate, distance of 57cm) and performing a feature-based attention task for liquid reward delivered by a custom-made valve system (Oemisch et al., 2019).

### 2.2. Data Analysis

Analysis of spiking and local field potential activity was done with custom MATLAB code (Mathworks, Natick, MA), utilizing functions from the open-source Fieldtrip toolbox (http://www.ru.nl/fcdonders/fieldtrip/).

### 2.3. Adaptive Spike-Artifact Removal (ASR)

We devised a novel method to remove spike-bleeding through artifacts from the local field potential recordings from the same recording channel data. This adaptive spike-artifact removal procedure, or *ASR*, has three main steps, delineated in **Figure 1**. These include (1) feature extraction, (2) decomposition and removal, and (3) reconstruction and integration. The main assumption underlying this technique is that the leakage of fast action potentials of a single neuron onto the LFP can be decomposed into a series of frequency components that can be robustly estimated. Following this estimation step, the spike-specific frequency components can be removed from the LFP, leaving a signal that is void of time-locked spike artifacts. An illustration of the underlying algorithm is provided for an example spike-LFP pair in **Figure 2**. Annotated programming code implementing the algorithm in Matlab and applying it to a test dataset is available as free open source software at www.accl.psy.vanderbilt.edu/resources/code/.

**Figure 1.**
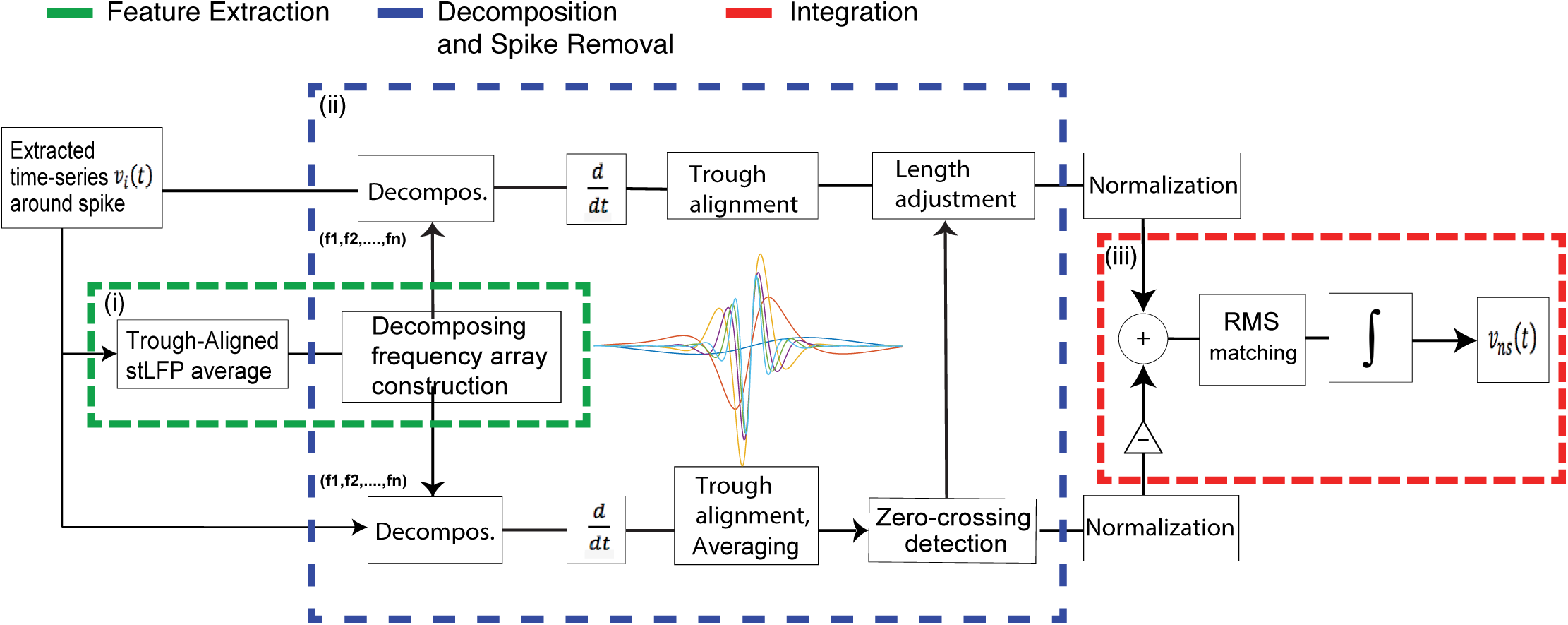
Adaptive-Spike-Removal consists of three main steps. i) feature extraction, ii) decomposition and removal, and iii) integration and reconstruction. In the first step (i, dashed green square), all 800-ms long trough aligned spike triggered LFPs (stLFP) are averaged. Then, the smallest frequency for decomposition is detected from the maximum peak in the derivatives of power spectral density of the average stLFPs and is used to construct a sequence of frequencies for the decomposition. In the second step (ii, dashed blue square), the spike triggered LFP for each individual spike is then decomposed into bands around the frequencies detected in step (i). In two parallel steps, first, the average of first derivative of trough aligned decomposed spike triggered LFPs is calculated. For each frequency component, the length of removal is determined based on the first zero crossing points of the decomposed stLFP to the left and right from the trough aligned center that covers peak values of one standard deviation above the mean of peak values. Then, each length-adjusted decomposed stLFP is normalized and subtracted from the normalized first derivative of the trough-aligned decomposed spike-triggered LFPs in the other parallel pathway and root mean square (RMS) of the after-subtraction signal is matched to the local signal RMS value. After the removal procedure on all frequency bands and all trough-aligned decomposed stLFP’s, the spike free signal (for the extracted duration around each spike) is reconstructed by summation across components and integration across time of all its after-removal first derivative of trough-aligned decomposed spike-triggered LFPs portions, using as starting value the value of the signal at the left hand side of the integration interval (iii, dashered square).

**Figure 2.**
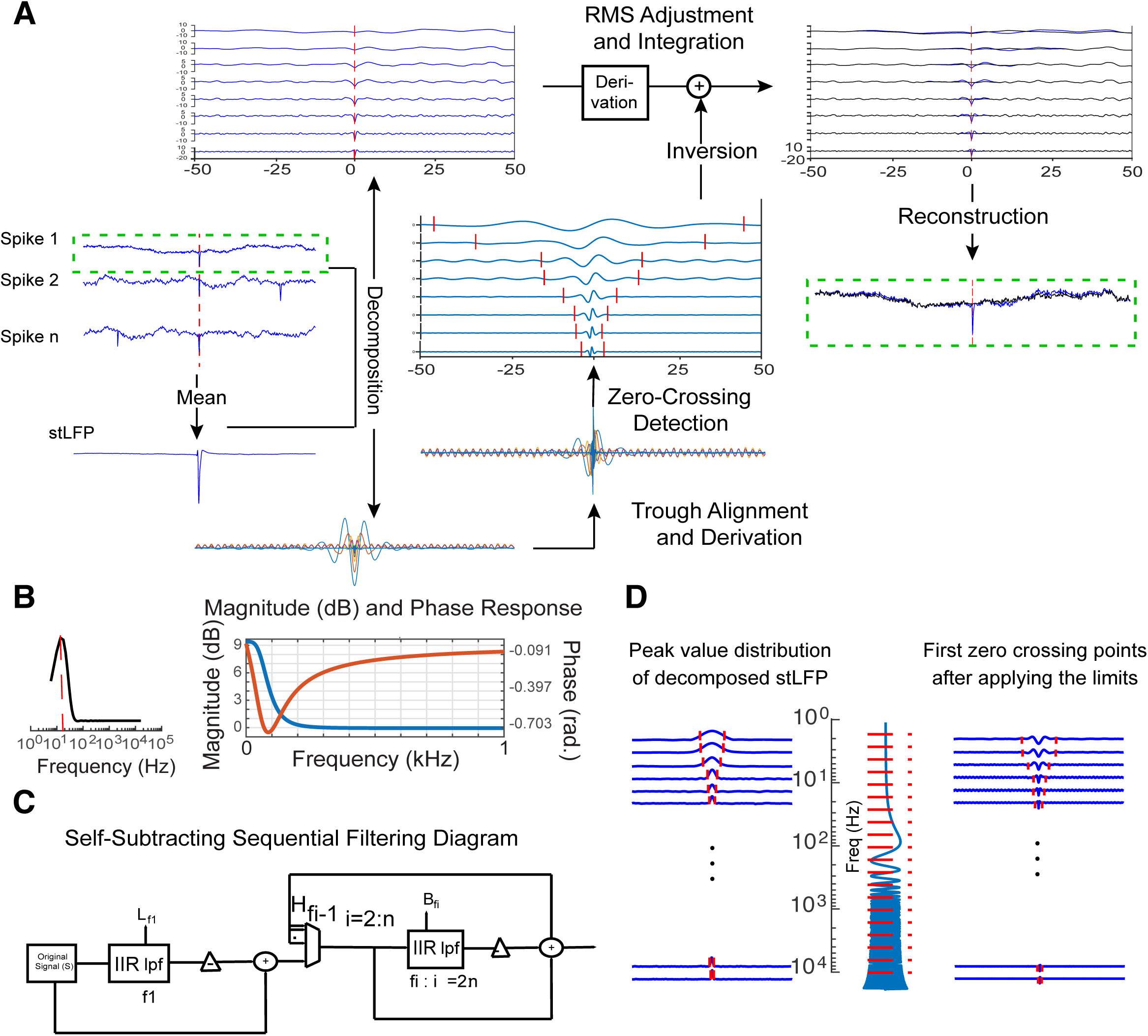
The spike removal algorithm applied to an example neuron. A. The spike-removal algorithm starts by calculating the average spike triggered LFP (stLFP). Then, after finding the center frequencies defining the bands for decomposition, each component of the spike triggered LFPs is averaged across spikes and the derivative is taken (d-stLFP). For each d-stLFP portion, the zero crossing points are detected (indicated by the red ticks) and after the length adjustment, each d-stLFP portion is trough aligned to the spike under consideration and removed from its corresponding portion of first derivative of decomposed stLFPs (to make the Figure simpler, the procedure is shown only on one stLFP, and for a subset of the decomposed portions). After removing each d-stLFP frequency components, the signal RMS value was equalized to the (non-spike) signal prior to the removal to preserve the signal mean power value. At the final step, all spike-free components are integrated, and their sum reconstructs the spike-free LFP. B. Frequency and magnitude response of a 50 Hz, low-pass filter. C. The block diagram of the sequential low-pass filtering design of the ASR method. Each level of filtering is performed on the subtraction of the previous low-pass filtered signal from the unfiltered signal. At the end the summation of all portions will reconstruct the original signal. D. For all decomposed spike triggered LFP’s, the distribution of absolute peak values is found. Then two initializing points before and after the spike trough were detected which define the edges of a time window with more than one standard deviation from the mean of average absolute peak values. For each band-passed portion of the signal, the first zero crossings were detected after these initializing points (red lines on the right panel).

#### 2.3.1. Feature extraction

In the first step of the ASR method we extract the frequency components of the spike-triggered LFP (stLFP) that are introduced by the spike event itself. To achieve this, we first calculated the stLFP in a ±400 ms window around the spike trough, averaged across all extracted spikes. For each individual spike event, the stLFP can be decomposed in its Fourier components (eq. 1.1), in which ω_0_ = 2πf_n_ is the angular velocity, a_n_ the Fourier amplitude of each frequency component, t time, and *k* the number of Fourier frequency components included:

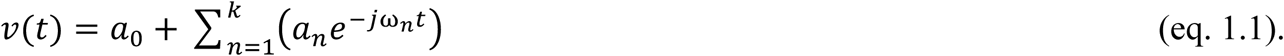

Each individual spike time series can be separated into two time-series v(t) = v_ns_(t) + v_s_(t), a spike triggered time-series:

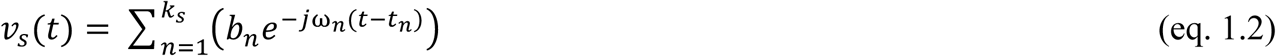

and a spike-free time-series.

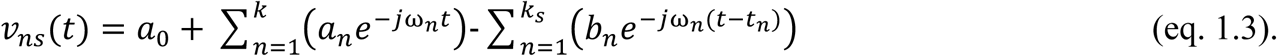

In the equations above, k_s_ denotes the number of frequency components of the spike time series, b_n_ is the scaling factor for the spike frequency time series, and t_n_ denotes the time lag for each frequency component from the spike trough. The relevant Fourier components were determined in a data driven fashion (see below). However, for each neuron, we assume that the spike waveforms contain components with the same frequency, but their amplitude and phase may be shifted from spike to spike.

In previous methods (Pesaran et al., 2002; Zanos et al., 2012; Zanos et al., 2011), all spikes from a given neuron were assumed to have the same temporal profile of their action potentials. However, the temporal profile of action potentials is not fully stationary and can vary depending on synaptic activity and axonal conductance (Bakkum et al., 2013). To accommodate these insights, we decomposed the stLFP of each individual spike to a series of frequencies with a distance of bandwidth (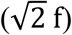) of a custom designed filter (illustrated in **Figure 2A**). However, for different neurons the starting frequency used in the decomposition is given by the smallest frequency that shows a higher power spectral density compared with its adjacent lower and higher frequencies. To find the lowest frequency of decomposition we computed the power spectral density (PSD) of the average stLFP. Then, we calculated the frequency normalized PSD and found the largest peak in a frequency range of 2-200 Hz. We then built an array of frequencies, sequentially separated by distances of 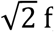, (which is the band limit of the band pass filter used), starting from the largest peak (**Figure 2A**).

To design a band-pass filter without distortion of the signal, we first designed a low-pass Butterworth filter (with an upper/lower band limit of ∼3dB, the magnitude and the phase response of a 50Hz low pass filter designed with this method is shown in **Figure 2B**). Then, we subtracted the low-pass filtered signal from the signal before filtering. The after-subtraction signal was again used for the next level of filtering (**Figure 2C**). In summary, each band-pass filter was built from two subsequent low-pass filters. By using this filter design, we were able to recover the original signal with adding all band-pass filtered signals, the first low-pass filtered signal, and the last high-pass filtered signal. The fact that each removal step is independent of others ensures that this method prevents any phase distortion when we sum up all filtered portion of the signal. Each filter was centered at the frequency peaks with an upper/lower band limit of 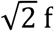, (∼3dB).

#### 2.3.2. Decomposition and removal

After the first feature extraction step we decomposed the signal further and then removed the spike components (**Figure 1**). For each of the bandpass filtered portions of the signal we calculated the average of the first derivative (differentials, eq. 2.1) of the spike triggered LFPs (d-stLFP). Under the assumption that non-spike components contribute to the averaged stLFP at random phases relative to the spike time, they add up to a constant, which is zero in our case, when averaged across a sufficient number of spike waveforms,

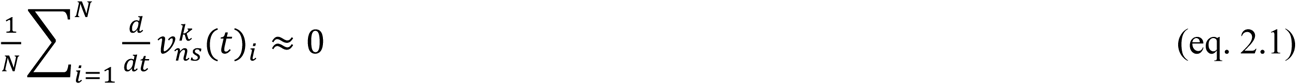

so that the average stLFP only contains spike-locked components:

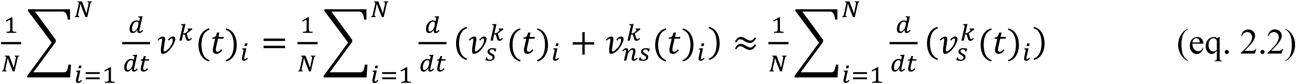

In Eq. 2.1 - 2.2, k denotes the *k*^th^ component of the spike time series, and i denotes the i_th_ spike. Then, the average of each d-stLFP component for all spikes were computed and normalized to its absolute trough value (in a complete trough aligned frequency cycle around the trough of the spike) and then subtracted from its correspondent d-stLFP component of each individual spike event normalized to its absolute trough value (in the same trough aligned complete frequency cycle).

To find the length of removal, we set two limits before and after the spike trough to search for the nearest zero-crossing of each averaged d-stLFP component after these two initial points. This initialization step of setting the minimum limit of zero-crossing points was performed in order to, first, find the extent of the artifact in each frequency band, then, considering the relative contribution of artifact before and after the spike trough. These limits cover a time window with d-stLFP absolute peak values of one standard deviation above the mean of d-stLFP peak distribution around each spike time in the corresponding frequency bands (**Figure 2D**). Because both the average and individual signal are normalized, the non-spike signal size after the subtraction does not match the actual signal size. We correct for this by considering the difference between the root mean square (RMS) of the total signal of a given spike before subtraction minus the RMS of the spike-locked signal as the correct size. Then we renormalize the subtracted signal so that its RMS becomes equal to the difference of the original signal and spike component RMS.

#### 2.3.3. Integration

After repeating the feature extraction and decomposition procedures (eq. 2.3) for all spikes and frequency components, the spike-free LFP was reconstructed by integrating the result, adding to the initial value (eq. 2.4).

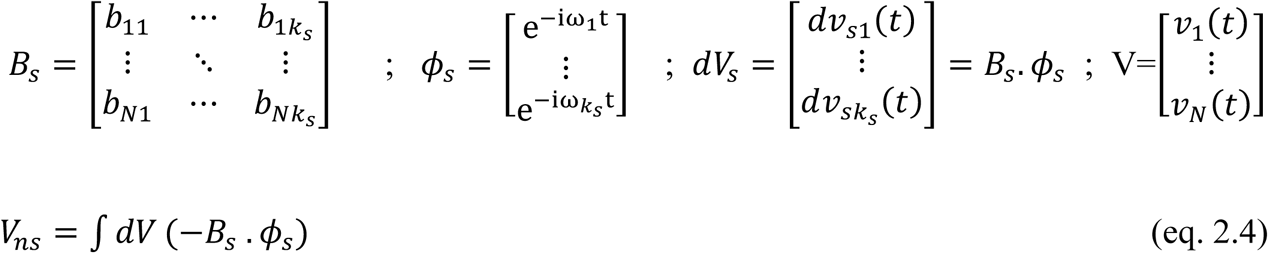

In the equation above, *k*_*s*_ denotes the *k*_*th*_ spike component, *N* denotes the N_th_ spike event. *dV*, is the derivative of the original spikes, *B*_*s*_ and ϕ_*s*_ are the computed approximations of all spike constant and phase matrices, and *V*_*ns*_ is the final output which is spike-free signal. Note that in the algorithm we do not explicitly calculate the parameters *B*_*s*_ and *ϕ*_*s*_, rather we represent them by the band-passed components in the decomposition. **Figure 2A** illustrates the removal algorithm on an example spike-LFP pair.

### 2.4. Data Simulation

In order to test, quantify and compare the ASR method using ground truth data, we simulated a LFP time-series and added spikes to the LFP (**Figure 3**). To make the LFP time series, we created a time-series that was the sum of two second-order autoregressive processes, with peak frequencies 30 and 50 Hz, respectively, together with a narrow band 10 Hz oscillatory signal that was created as follows. We simulated a phase time series by integrating across time a constant frequency of 10 Hz (varied from 10 to 80 Hz) plus a small normally distributed random component, independent across time steps. The third component was then obtained as the cosine of this phase times a scaling factor. This phase was used also used to align the spike times. A spike was placed in the signal when the strength of the target oscillations to which it needs to be locked exceeds a certain strength, the spike time is than given by a normal distribution around the target phase. We achieved this by translating these constraints into a time-varying Poisson process with as integrated sum the number of spikes we desired (this is an approximation as locking of spike times to input features contains non-Poisson statistics, see (Toups et al., 2011, 2012)). Spikes are modeled using an asymmetric triphasic template modeled as a Gabor function, a Gaussian multiplied by a cosine offset by a phase from the center of the Gaussian, but with variable voltage amplitudes to also mimic the potential axonic conductance variations explained before (**Figure 3A**) (Gerstner et al., 2014).

**Figure 3.**
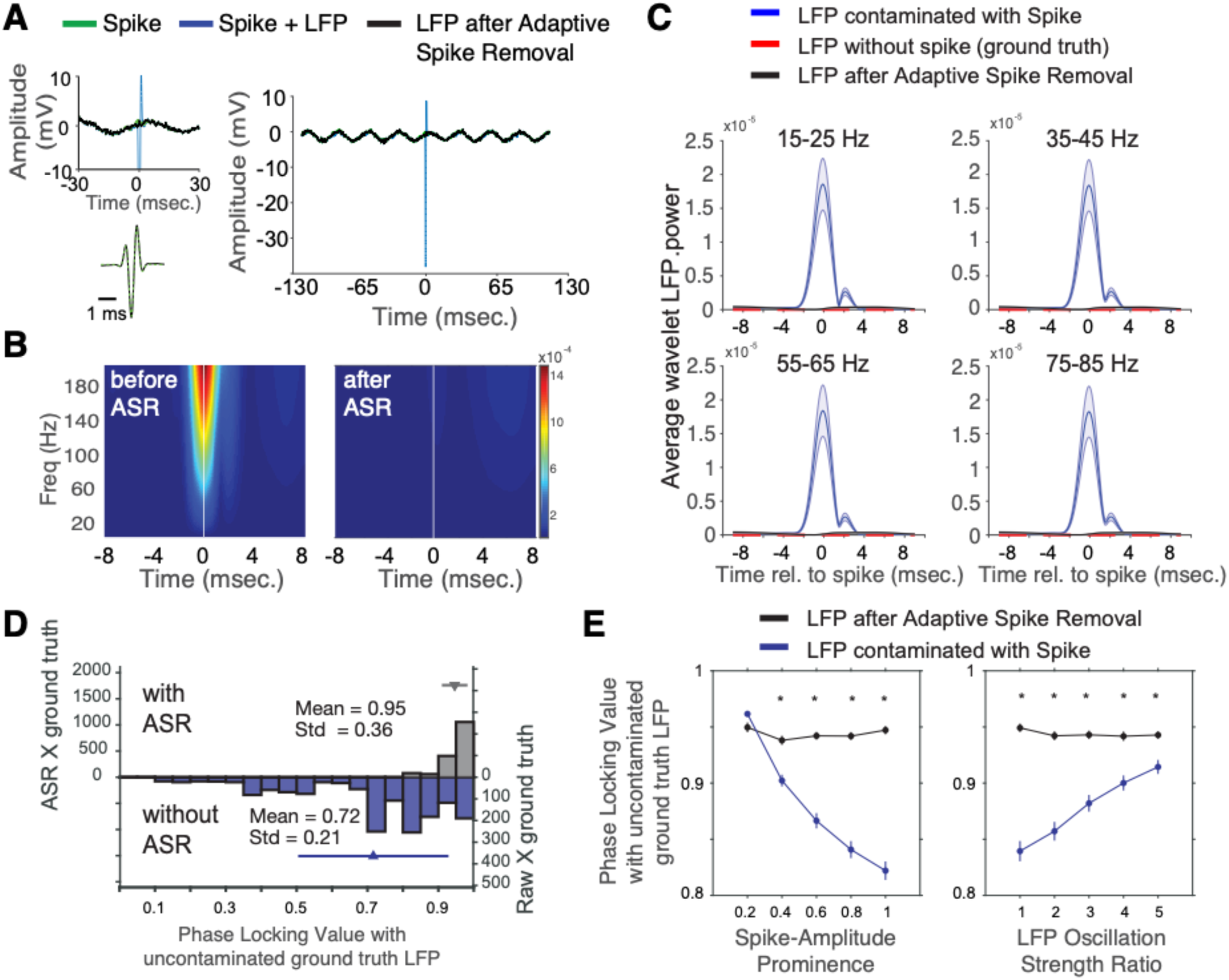
Applying Adaptive Spike Removal (ASR) on synthetic ground truth data. A. Spike-triggered LFP average (stLFP) for synthetic ground truth data (green), for ASR (in black), and spike-contaminated data (blue) is shown and the spike extracted with the ASR method is shown on the bottom. No edge artifact is introduced to the signal. B. A time frequency analysis of spike triggered power is shown for the same simulated unit. The spike-related power transient is effectively removed by the ASR method. C. The wavelet analysis of spike-triggered average power in four frequency bands for the ground truth data with spike (blue), without spike (red), and when the spike was removed using ASR (black). The results are the average across 100 simulated units showing that the ASR method removed the spike artifact without changing the power of ground truth LFP (the black line falls within the range of the standard error of the red line). D. Distribution of phase locking values between ground truth LFP (uncontaminated by spike) and with LFP that was contaminated by spikes (blue bars), and with LFP after adaptive spike removal of contaminating spikes (grey bars). E. The phase locking value computed as in *E*, but with increasing spike amplitude relative to the LFP amplitude (*left*), and with increasing oscillation amplitude at constant spike amplitude (*right*). ASR lead to a slight attenuation of the PLV (a perfect match would result in a PLV of one), but otherwise is unaffected by spike prominence or LFP oscillation amplitude.

To simulate low frequency spike-related transients, we added three cycles from 20 Hz, 55 Hz, and 85 Hz frequency components to locked to the spike time with phases that have a modest standard deviation. We then used the spike-free LFP as the ground-truth signal and applied the ASR method to the spike-injected LFP. We repeated this procedure for 100 simulated cells with varying firing rates (5-50 spk/sec), frequency of phase coherency (10 Hz to 80 Hz) and spike-to-LFP amplitude ratios (10 to 50). To evaluate the spike removal, we computed the stLFP for all three time-series, as well as the wavelet analysis of the spike triggered power spectrum in a ±500 ms window around the spike trough (**Figure 3B**). To quantify the difference between the ASR results, the spike contaminated LFP, and the ground-truth signal, a wavelet analysis was applied to the difference of each time-series from the ground-truth signal at the constructed spike times (**Figure 3C**).

### 2.5. Spike Removal using the Bayesian Removal

A prior study proposed a Bayesian spike removal approach on LFP data from visual cortex (Zanos et al., 2011). This Bayesian spike removal method reconstructs the wideband signal around the time of the spike by summing a set of low-frequency oscillations, the stereotyped spike waveform around the spike time and the noise. The cleaned spike-removed low-frequency LFP is then estimated using Bayesian inference. To compare this Bayesian approach to our adaptive spike removal method, we used the analysis code and datasets provided by Zanos et al. (2011) and applied both the Bayesian and the ASR method to their datasets (**Figure 5**).

### 2.6. Spike Removal using the Average Field with interpolation

In addition to the adaptive ASR removal method we also tested the effects of removing the average spike artifact from the stLFP (Zanos et al., 2012). This average removal method has been used before e.g. in datasets in which the wideband data were not available in the high frequency range at which the action potential has highest power (Ardid et al., 2015; Zanos et al., 2012). For the average removal method a duration of 5 ms was extracted from all trough-aligned spikes (from 2 ms before to 3 ms after the trough) for each neuron and their average was removed from each individual spike in the wideband data to eliminate spurious spikes from the LFP (the maximum length of removal to not remove more than a complete cycle up to 200 Hz is 5 ms).

### 2.7. Quantification of spike-LFP synchronization

We used the fieldtrip toolbox for Fourier analysis of the LFP. The raw signal was resampled to a 1000 Hz sampling rate. The Fourier transform was performed on 5 complete frequency cycles using an adaptive window around each spike (two and a half cycles before and two and a half cycles after the spike). Then, the pairwise phase consistency (PPC) was computed to measure spike-LFP synchronization (Vinck et al., 2012; Vinck et al., 2010).

We determined whether the spike-LFP synchronization (PPC) spectrum contained peaks that signify statistically significant, frequency-specific phase-consistent spiking using four criteria similar to (Ardid et al., 2015). First, peaks had to have a Rayleigh test significance of p<0.05 to reflect statistically that the phase distribution was not homogeneous. Second, each peak had to have a PPC value greater than 0.005. Third, each peak had to have a minimum local prominence of 0.0025 from its two neighbor local minima, which rejected locally noisy and possibly spurious PPC peaks. Fourth, peak values had to be greater than 25% of the PPC range.

### 2.8. Phase and power comparison

The spike triggered average LFP power was calculated with a moving window of ±100 ms around each spike every 5 msec. We also measured mean phase and phase distribution of spike triggered LFP from the Fourier transform using standard circular statistics (Berens, 2009).

## 3. Results

### 3.1.1. Adaptive Spike Removal (ASR) on simulated spike-LFP data

We first tested the Adaptive Spike Removal (ASR) method on synthetic data with known ground-truth spike waveform shape and known frequency and noise components of the LFP. In simulations we ensured that spikes injected to the simulated LFP introduced the typical spike aligned bleed through into the lower frequency band of the LFP (**Figure 3A,B**). Applying ASR to the spike-triggered LFP of 100 simulated neurons with varying spike amplitude and firing rates efficiently removed the spike artifact, leaving no discernible power spectral component of the spike in the cleaned LFP data. We quantified the removal quality by calculating the wavelet power spectral density of the spike-triggered LFP before and after ASR in four lower frequencies (15-25 Hz, 35-45 Hz, 55-65 Hz, and 75-85 Hz). The results from this analysis illustrate that the ASR method retrieved the original uncontaminated LFP with no apparent residuals from the artificially injected spikes (**Figure 3C**).

In order to test how robust the ASR method removes spikes with different amplitudes relative to the local field potential (i.e. different signal-to-noise ratios) we simulated 15000 ground truth datasets and calculated the phase locking value between the ground truth LFP with the ground truth LFP that was injected with spikes at different oscillation frequency (between 20Hz and 180 Hz), and different amplitudes (trough amplitude relative to the LFP amplitude ratio’s from 0.2 to 1). First, we found that the phase locking value of ASR treated data with the uncontaminated ground truth data was close to one, while ground truth and contaminated data showed reduced PLV (**Figure 3D**, significantly higher PLV for ASR treated data, paired t-test P<.0001). Next, we found that ASR had a constant small effect of attenuating the similarity with the ground truth data with a phase locking value with ground truth data of ∼0.95, but that this attenuation was unchanged with increasing spike amplitudes (**Figure 3E**, left). In contrast, without ASR, the PLV systematically decreased with increased spike amplitudes. Similarly, the ASR was unaffected by varying the oscillation amplitude in the LFP, while increased oscillation amplitudes reduced the PLV for the spike-contaminated data (**Figure 3E**, right).

### 3.1.2. ASR on simulated spike-LFP data with high, 160-200 Hz oscillatory LFP components

In some neural systems like the hippocampal formation, activity in the high gamma, epsilon, or ripple band (e.g. 80-200 Hz) carries important functional signals and is prone to be contaminated by spikes (Buzsaki and Silva, 2012; Leonard and Hoffman, 2017; Leonard et al., 2015; Schomburg et al., 2014). We therefore tested how the ASR performs for spikes contaminating high LFP frequencies and quantified in simulations at which oscillation frequency the method fails to reliably remove spike from the LFP. We simulated data with oscillation frequency of 160, 180, and 200 Hz while varying the power peak frequency of injected spikes from 500 Hz to 300 Hz (**Figure 4A**). Applying the ASR algorithm to these data showed that the ASR method works as long as the ratio of the frequency with peak spike power to the frequency with the LFP power of interest is larger than √2. We quantify this by showing constant high phase locking values for LFP data as long as the spike peak power is 350Hz or higher (**Figure 4B**). However, applying ASR to data with high frequency oscillations will lead to a power attenuation of the wideband data around the time of the spike depending on the degree to which the frequencies with spike power overlap the frequency of interest in the LFP data (**Figure 4C**). This attenuation might be conceived of a limitation of the ASR method, but it stems from the successful removal of spike related frequency components, which is the desired goal of the ASR algorithm.

**Figure 4.**
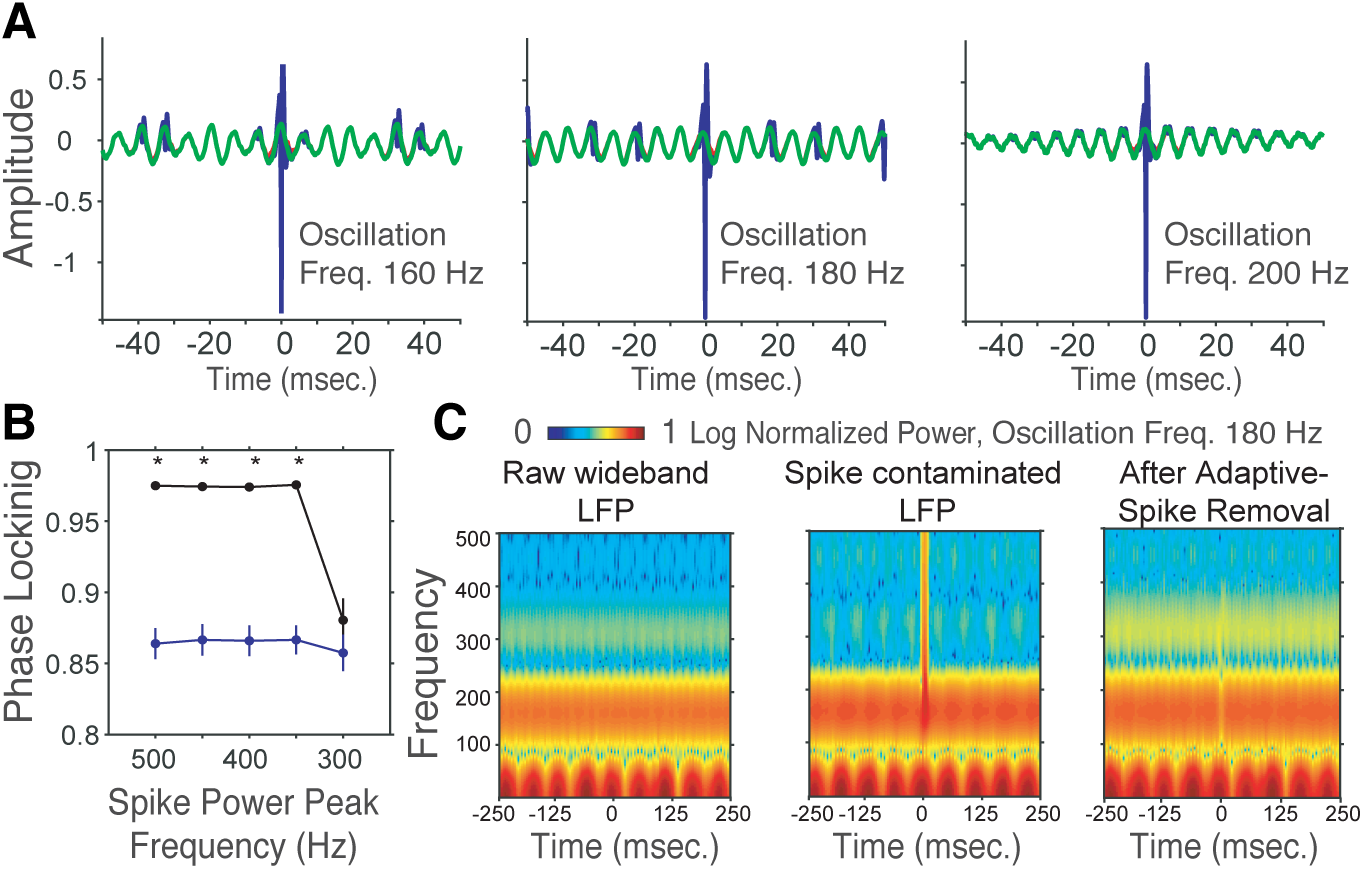
Constraints for Adaptive Spike Removal at High Oscillation Frequencies. A. Illustration of simulated wideband data with oscillation peak frequencies at 160, 180, and 200 Hz (green) with added action potentials (blue). B. Phase locking value for the phase consistency between the ground truth (uncontaminated) LFP and the LFP with ASR (black), and between the uncontaminated versus contaminated ground truth. The x-axis varied the width of the action potential that contaminated the simulated data with spectral peak power of the spike ranging from 500 to 300 Hz. C. Time frequency spectra aligned to the time of the spike in the uncontaminated ground truth data (left), the contaminated ground truth with spike bleed through (middle), and after applying Adaptive Spike Removal (ASR) (right). ASR slightly attenuates the power around the time of the removed spike at these high frequency because of overlap in spike and oscillation frequency components.

### 3.2. Comparison of ASR and Bayesian spike removal

Prior studies have suggested that a Bayesian approach is an efficient method to remove spike-related transients at low LFP frequencies (Zanos et al., 2011). We therefore evaluated the Bayesian spike removal method with the ASR method by applying both approaches to the data provided by (Zanos et al., 2011). We found that the performance of ASR is superior to that of the Bayesian approach, confirming our initial impression that the Bayesian approach does not prevent phase distortions (also noted by (Pesaran et al., 2018))(**Figure 5**). At high ≥100Hz frequencies the Bayesian approach does not remove spike artifacts, while ASR effectively removes spike-specific components from the LFP evident in the spike triggered LFPs (**Figure 5A**) and in the wavelet decomposed spike-triggered averages (**Figure 5B**). For lower (<100 Hz) LFP frequencies ASR removes spike-related power transients thoroughly, without noticeably changing the power in low frequencies elsewhere from the spike time origin, while the Bayesian method removes LFP components at low frequencies even if they have no apparent relation to the spike (**Figure 5C**).

**Figure 5.**
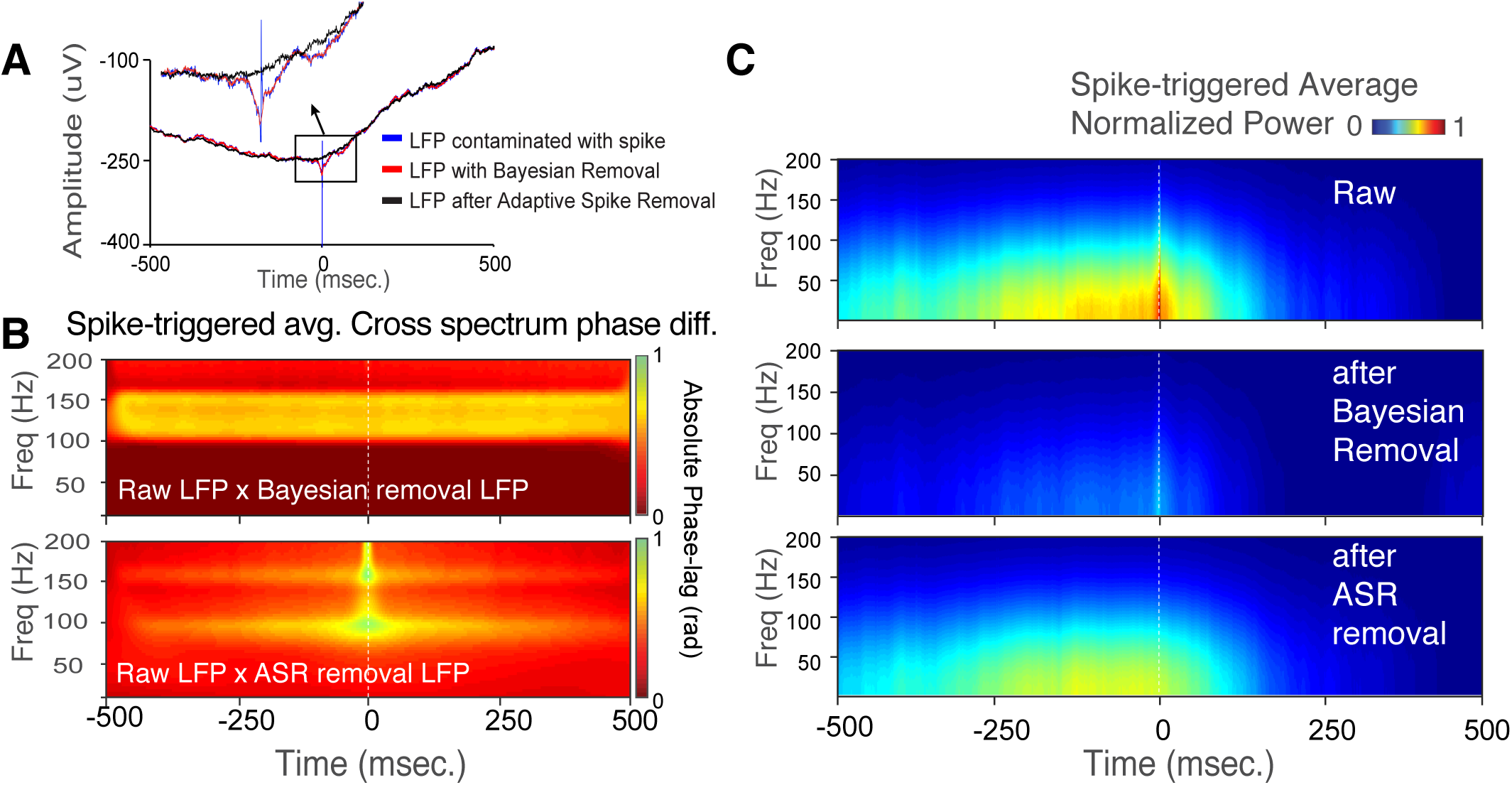
Bayesian and ASR comparison. A. Spike-triggered LFP average illustrates the improvement of ASR (black), over the Bayesian (red) approach to remove spike-related transient (blue) in low frequencies. B. The spike-triggered cross-spectral phase delay of the raw signal and the signal after Bayesian spike removal (top panel) and ASR (bottom panel). The ASR method has a higher specificity for removing spike-related transients from the LFP centered at the spike time, while the ASR does not affect the phase away from the spike time. C. Temporally resolved spike-triggered LFP power is shown for raw signal (top panel), after applying Bayesian spike removal (middle panel) and ASR (bottom panel). While Bayesian removal method has still artifactual leftovers in the spike-related frequency range, the ASR method is able to remove spike-related power transients without noticeably changing the power in low frequencies elsewhere from the spike origin.

### 3.3. Using ASR to elucidate beta frequency spike-LFP relationships

The results so far illustrate that the ASR method is an effective way to clean LFP data from spike-specific artifacts across a broad frequency range that encompasses the beta, low gamma and high gamma frequency range. We thus tested whether ASR can be used to clean LFP data from spiking activity recorded at the same recording channel, allowing quantification of local phase synchronization of the spike timing to LFP. To test this, we recorded wideband LFP data in nonhuman primate anterior striatum and calculated spike-triggered average wideband activity aligned to the trough of spikes from single neurons whose spike times were identified using a spike-sorting procedure (Oemisch et al., 2019). Across multiple examples of raw spike triggered LFP data we observed the expected, apparent spike-bleed through artifact that prevented discerning a veridical spike-phase relationship to the LFP for frequencies as low as 20 Hz and upwards (**Figure 6**). This is illustrated for an example neuron in **Figure 6A**. This neuron showed enhanced spike-LFP synchronization starting at ∼20 Hz. But using the raw LFP data did not allow inferring whether this ∼20-40 Hz phase synchronization was real, or artifactually contaminated by spike-bleed through because the synchronization peak in the beta band is overlapping with the frequency range that is obviously affected by spike-bleed through evident in the continuous rise of spike-LFP synchrony from 40Hz onwards and beyond >200Hz (**Figure 6A**, *middle panels*). In contrast, applying ASR removed the spike artifact effectively as evident in the spike triggered LFP (**Figure 6A**, *left panels*), the wavelet decomposed spike triggered LFP and the phase spectrum (**Figure 6A**, *right panels*). Consequently, following ASR the neural spike times show a clear 20-40 Hz spike-LFP phase synchronization peak. Notably, for frequencies below 20Hz the spike-LFP synchronization estimates are similar irrespective of whether spike removal is done with ASR or with average spike removal confirming a prior study’s report (Ardid et al., 2015).

**Figure 6.**
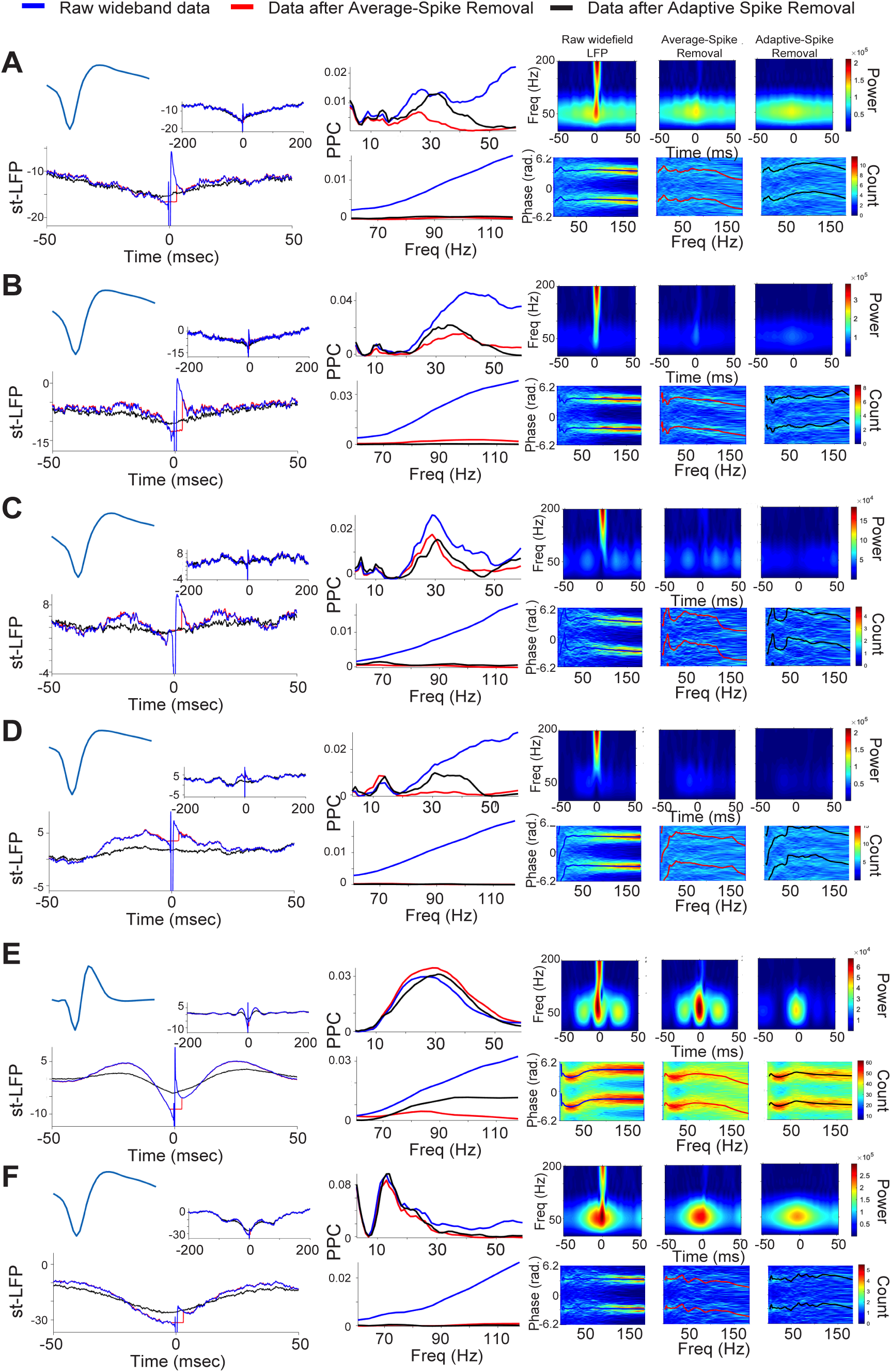
Example neurons documenting spike-LFP relationships of signals recorded from the same tungsten electrode without spike removal (raw), with Average-Spike Removal (red) and ASR (black). A-F. The left top panel shows the average action potential waveform of the recorded neuron. The spike-triggered averages are shown for ± 50 ms time lag and as an inset for ±200 ms. The middle column shows the spike-LFP phase synchronization measured as the pairwise phase consistency in the 2-60 Hz range (upper panel) and 60-120 Hz (lower panel). The color panels on the right shows the spike-triggered wavelet decomposition (upper panels) and the 3D phase histogram (lower panels) for (from leftmost to rightmost panels) the raw LFP, after Average-Removal, and after ASR. Examples in A-F document how ASR effectively removes the spike artifact and uncovers spike-LFP phase synchronization in the 20-35 Hz beta frequency band.

The described spike-LFP pair is a representative example. **Figure 6B-E** illustrate four additional examples for which ASR uncovered narrow band spike-LFP phase synchronization in the 20-35 Hz beta frequency range that would have been unrecognized without spike removal. In contrast to this 20-35 Hz range for which ASR can be a necessary method to uncover spike-LFP synchrony, spike-LFP locking in a 15-20 Hz band (or lower) does not depend on spike removal. **Figure 6F** illustrates such a case with spike-LFP synchronization in the low beta band that was evident independently of whether the spike was removed from the LFP or not.

Another result of the analysis of example neurons in **Figure 6** is that ASR is effective for neurons with different action potential shapes (shown as insets in each panel). Action potentials of neurons in the striatum range from narrow to broad waveforms, exemplified in the examples in **Figure 6E** (narrow) and **Figure 6A** (broad) (Berke et al., 2004; Lansink et al., 2010). The ASR approach is tailored to adjust the removal adaptively according to the specific action potential waveform of the neurons. Thus, ASR allowed to quantify beta band spike-LFP relationships that would be difficult to access without effective, adaptive spike artifact removal. This conclusion is supported also by the average spike-LFP synchronization across the population of all recorded neurons, which revealed a particularly increased gain in observing ∼20-45 Hz synchronization with ASR compared to non-adaptive spike removal methods (**Figure 7**).

**Figure 7.**
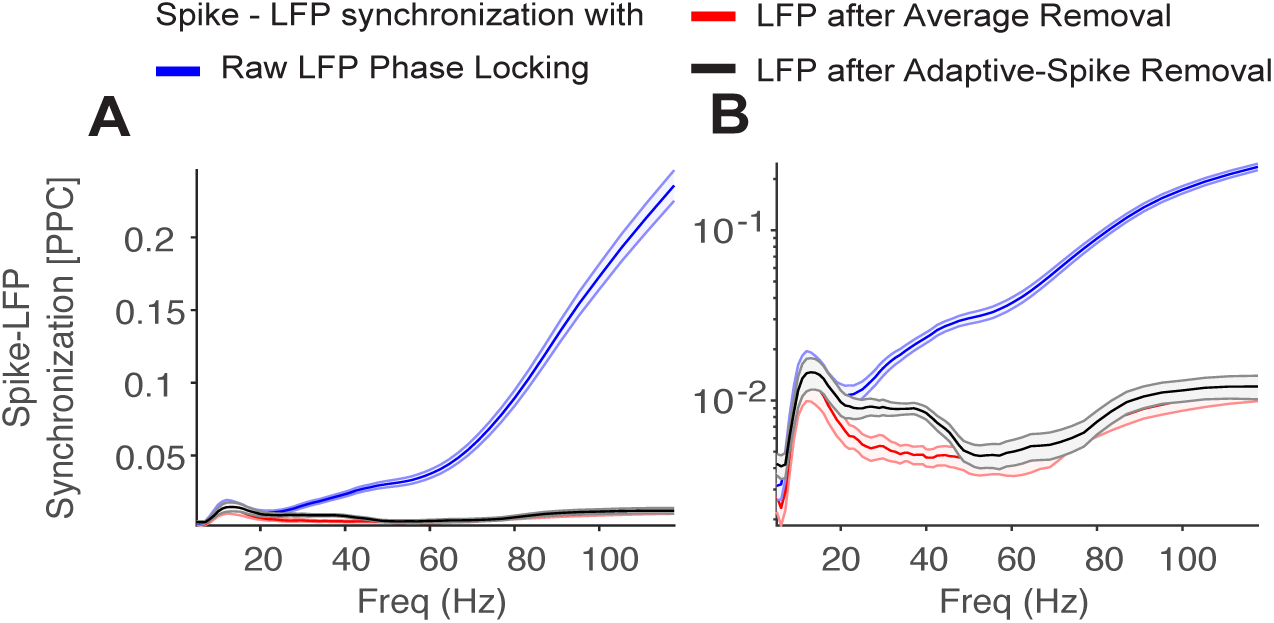
Average spike-LFP synchronization across all striatal recorded neurons. A and B shows the same data with different y-scaling, illustrating that with ASR (black), stronger average beta and low gamma band spike-LFP synchronization is evident that would otherwise be hidden by the spike-bleed through artifact in the raw LFP data (blue), and is not detected by the Average-Removal approach. Error bars denote standard error.

### 3.4. Identifying gamma frequency spike-LFP relationships using ASR

There has been no method available so far to robustly estimate gamma frequency phase synchronization of spike times to the local, same-channel LFP, because the spike timing relationship to gamma band LFP fluctuations are difficult to quantify without removing spike-bleed through (Ray, 2015). We thus quantified how the ASR approach worked for spike triggered LFP’s that showed apparent oscillatory sidelobes at time lags indicative of ∼35-45 Hz low gamma band oscillations (see examples in **Figure 8**). Applying ASR successfully removed the spike artifacts for these cases and allowed quantifying prominent gamma band spike-LFP synchronization (**Figure 8A-F**).

**Figure 8.**
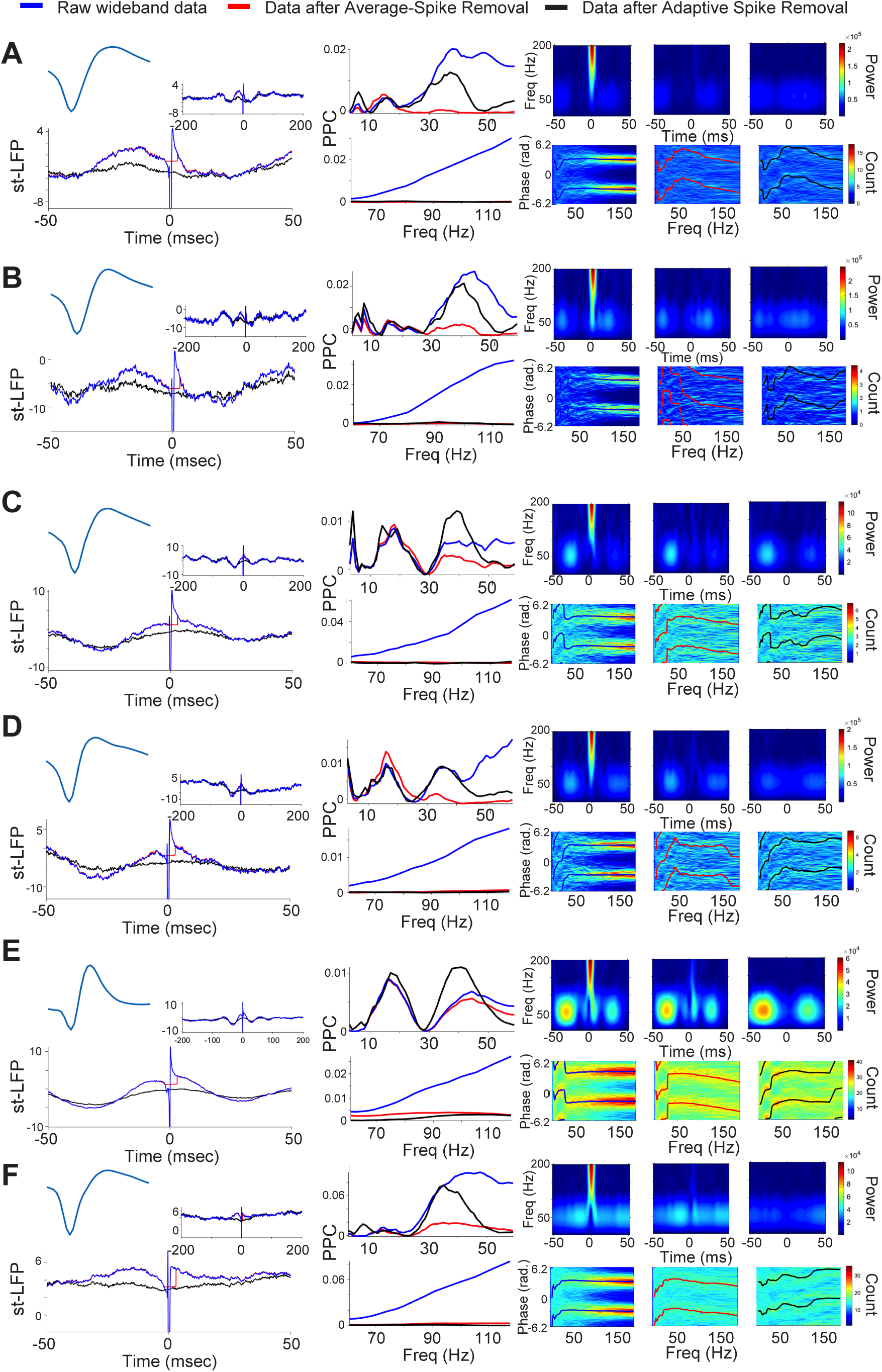
Example neurons documenting spike-LFP synchronization in the 30-45 Hz gamma frequency after ASR spike removal. Same format as Figure 5, for spike-LFP pairs that showed apparent spike-locked gamma oscillations in the spike triggered LFP average. Using ASR this gamma band synchronization becomes quantifiable and evident in separate peaks of the PPC spectrum.

To summarize the results of ASR for the whole neural population, we estimated spike-LFP synchronization for all 293 striatal single neurons recorded with wideband LFP (Oemisch et al., 2019). For each spike-LFP pair we calculated the pairwise phase consistency of spikes to the raw LFP and to the LFP after removal of the spike artifact using the ASR method. We found that ASR significantly increased the number of spike-LFP pairs with significant and robust peaks in the PPC spectrum (**Figure 9**). We found that 164 (56%) of spike-LFP pairs had peaks in the phase synchronization spectra in the raw, contaminated data (**Figure 9A**), 202 (69%) of spike-LFP pairs had peaks when applying the average removal methods from Zanos et al., (2012) (**Figure 9B**), and 250 (85%) of spike-LFP pairs had apparent synchrony peaks with ASR (**Figure 9C**). Thus, ASR allowed detecting significant phase synchrony in 29% of neurons whose spike-LFP synchrony was too contaminated otherwise (Z test on proportional difference p<0.001 for raw vs ASR, and p=0.001 for average removal method vs ASR). The majority of these additional spike-LFP synchronous cases showed phase synchronization in the beta and low gamma bands. In summary, ASR allowed to uncover spike-LFP relationships that would not have been available with other spike-removal methods.

**Figure 9.**
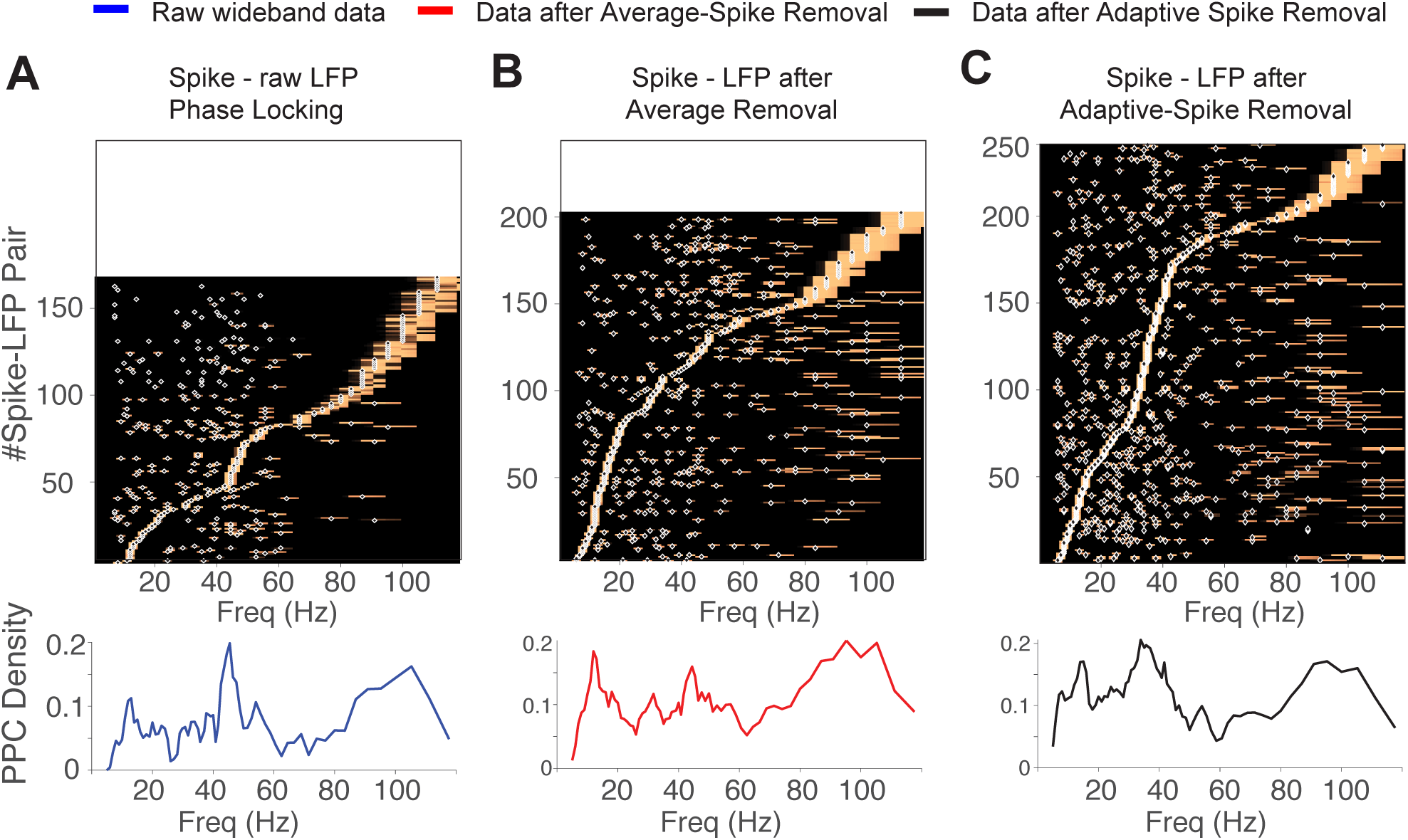
Distribution of significant spike-LFP synchronization peaks across the striatal neuron population. The top row shows spike-LFP synchronization normalized and ordered by the maximal peak value for each spike-LFP pair (y-axis) that showed at least one significant peak in the PPC spectrum. There were fewer significant peaks for spike-LFP pairs for the raw contaminated LFP (A) than after Average-Removal (B) and ASR (C). The bottom panels show the peak densities of spike-LFP synchronization calculated for each condition. The results illustrate that ASR uncovers more spike-LFP pairs with a significant beta and gamma band synchronization. The bottom row shows the density of peaks (y-axis) across frequencies (x-axis).

## 4. Discussion

We showed how a method that adaptively estimates the frequency components of individual action potentials of a neuron is capable to effectively remove the spike artifact that would otherwise bleed-through into the lower frequency range (<100 Hz) of LFPs. The described adaptive spike removal approach effectively extracts ground-truth spike action potential waveforms injected into synthetic wideband LFP irrespective of the amplitudes of the spike artefact. We then showed that the novel approach surpasses the existing Bayesian approach for removing the estimated average waveforms from local field potentials. Applying adaptive spike-artifact removal to neurons and local fields recorded from the same electrode in rhesus monkey striatum cleaned the data and made it possible to quantify spike triggered averages, spike-LFP phase consistency and spike-LFP phase spectra without apparent contamination of the action potential of the neuron itself. Most importantly is that the method uncovered narrow frequency phase synchronization at the beta and gamma band that would have remained contaminated otherwise. Taken together, these results suggest that the proposed adaptive spike-artifact removal method might become a new state-of-the art approach for cleaning wideband data from action potentials prior to same-channel or nearby-channel spike-LFP phase analysis.

### The algorithm’s advantage: Decomposing individual spike waveforms

One central reason for the success of the adaptive removal method is that it takes into account the variation of individual action potentials as opposed to estimating an average waveform. Previous approaches have either considered only the average waveform of a neuron to subtract its template (Pesaran et al., 2002; Zanos et al., 2012; Zanos et al., 2011), or removed data corresponding to the average time-course of a neuron’s action potential (Ardid et al., 2015; Galindo-Leon and Liu, 2010; Okun et al., 2010). Both of these approaches reduce the highest amplitude transient spike artifact, but they leave residual waveform components in the LFP that reduce the usability of the method (**Figure 5B,C**). Consistent with this caveat, we have previously shown that with these methods it is only possible to consider spike-LFP relationships to be void of phase distortions up to ∼25Hz (Ardid et al., 2015). A similar conclusion was drawn by (Pesaran et al., 2018) who recommended not to consider same-channel spike-LFP because of the residual leakage effects that makes them difficult to interpret with existing methods. Our results with the average (non-adaptive) removal methods agree with this conclusion.

In contrast to the average-waveform-based approaches the new adaptive spike-artifact removal approach decomposes not the average waveform, but all individual action potential waveforms separately. This approach acknowledges the variability in action potential shapes that have been well documented in in-vitro and rodent in-vivo work. For example, variability in axonal conductance influences the precise timing and causes variations in action potential amplitudes (Bakkum et al., 2013; Bucher, 2016). The variations in axonal conductance can be traced back to specific ion channel kinetics and densities (Ganguly et al., 2000), extracellular neurotransmitter and ion densities (Kocsis et al., 1983), presynaptic modulation by glia (Sasaki et al., 2011), or gap junction coupling and ephaptic interactions (Barr and Plonsey, 1992).

### A window into beta and gamma band synchronization within a 200 μm microcircuit

Our results show that the novel spike-artifact removal method is able to uncover phase synchronization of single neurons to 20-35 Hz beta and 35-50 Hz gamma band LFP activity recorded on the same electrode as the neuron itself (**Figure 9C**). We believe this finding illustrates that the method provides a novel window into studying single neuron-to-circuit interactions. Beta-band activity is ubiquitous in the non-human primate striatum with some studies reporting more than 90% of LFP power spectral densities showing beta rhythmic peaks (Amemori et al., 2018), and multi-unit activity synchronizing to beta rhythmic LFP measured from electrodes approximately 500 μm away from the location of the multi-unit spiking neurons themselves (Antzoulatos and Miller, 2014). To our knowledge there is no prominent example published in the nonhuman primate striatum with isolated single neurons spiking synchronized to the local circuit LFP in the gamma frequency band. Studies in the striatum of rodents report of gamma activity when analyzing spike and LFP from spatially separate tetrodes (Howe et al., 2011; Kalenscher et al., 2010; van der Meer et al., 2019; van der Meer et al., 2010), or have focused on LFP analysis without reporting local spike-LFP phase relationships (Catanese et al., 2016). Of note is that some forms of striatal gamma activity in the rodent do not have a local origin (Carmichael et al., 2017). However, the ASR method is agnostic to the source of possible oscillations or to the cell and membrane configurations giving rise to the LFP, and is focused just on removing the spike.

In this context it appears that our new ASR method was essential to have uncovered a larger population of single neurons in rhesus monkey striatum that phase synchronizes to the LFP at a narrow gamma band with peaks around 40 Hz (**Figure 8**). This gamma band spike-LFP phase synchrony would be expected for a small population of fast spiking interneurons (FSI’s) in the striatum whose intrinsic properties make them gamma rhythmic pacemakers (Belic et al., 2017). These FSIs are expected to have narrow spike waveforms, which we observed in some example spike-LFP pairs with gamma synchrony (**Figure 8E**) (Berke, 2009). However, beyond these narrow waveform neurons we see multiple other examples with broader action potential shapes and narrow gamma band peaks (**Figure 8A-D**). The morphological identity and the functional contributions of these neurons to the local circuit functioning in the striatum is to our knowledge unknown. The results from applying the adaptive spike-artifact removal are thus opening up a possibly new window into understanding the cell-to-circuit neuronal interactions.

### Comparison with other methods

Prior work has proposed spike-artifact removal using Bayesian methods. We show that these methods still leave residual phase distortions in low-passed filtered LFP (e.g. **Figure 5E**). To understand why these methods have been considered to provide a solution to the spike bleed-through challenge it might be important to consider in which neuronal circuits they were developed. The Bayesian removal was applied to our knowledge for visual cortex neurons and neither in (Zanos et al., 2011), nor in (Zanos et al., 2012) were their example neurons showing strong spike-LFP synchrony. In contrast, in our striatal recordings we found many examples of neurons with strong spike-LFP synchrony with neurons firing >1.2 more spikes at the preferred as opposed to the anti-preferred LFP phase. This prominent rhythmic phase synchrony is also visible in the oscillatory sidelobes of the spike triggered average LFP for many examples (**Figure 6, 8**). We have not seen comparably strong phase synchrony in prior studies evaluating artifact removal algorithms (Zanos et al., 2012; Zanos et al., 2011). We therefore believe that the strength or weaknesses of artifact removal approaches might depend on the presence of data that show clearly discernible spike-LFP phase relationships.

### Adaptive artifact removal for channels with spikes from multiple single neurons

One limitation of the current adaptive spike removal approach is that it best works on datasets with action potentials that are similar and hence likely originate from the same neuron. In many electrophysiological recordings, a single recording channel will have spikes from multiple different neurons (Buzsaki, 2004; Rossant et al., 2016). To practically address this situation the adaptive spike removal method could be applied sequentially on the same recorded channel to extract spikes from one neuron in the first run, spikes from a second neuron in a second run, etc. Such sequential adaptive spike removal could also prevent spike bleed though artefacts from spikes of neurons that co-activate with another neuron’s spikes. An alternative to this sequential cleaning of the wideband data is to perform the adaptive spike removal only once for each targeted action potential (neuron) type and repeating it using separate copies of the wideband data for estimated its spike-LFP phase synchrony for each unique neuron. However, we could imagine that future versions of the ASR can be tailored to clean wideband data from multiple variable action potentials if desired.

### Adaptive artifact removal for low frequency components of the spiking effect on the LFP

The good performance of our adaptive spike-artifact removal method is likely also be due to considering a wider temporal dynamics to contribute to the fast action potential than the brief 1-5 ms sharp transient spike itself. Prior biophysical modeling studies have shown that while spike-locked synaptic effects on the LFP rises sharply, they decay over longer ≥ 10 ms time windows than the duration of action potential repolarization (Schomburg et al., 2012). Furthermore, it is known that Na+ spikes of neurons working coherently, or in an asynchronous regime have afterpotentials that can last 2-20 ms (Fernandez et al., 2005), and spikes can trigger NMDA and Ca+ plateau potentials that are phase-locked to the spike time and prevail in the LFP (Buzsaki et al., 2012; Watson et al., 2018). These influences will account to different degrees for the slow, non-transient modulation that is visible in many raw spike-LFP triggered spectra. The proposed ASR addresses these slow, low frequency contributions to the LFP.

In summary, we propose an adaptive spike artifact removal approach and show that it outperforms existing approaches by addressing major limitations of these prior methods.

- ASR is adaptive to the center of the leakage frequency bands of individual neurons. This *frequency adaptive feature* prevents removing non-spike components from the signal and makes the algorithm flexibly applicable to widely varying spiking types including bursting neurons and fast spiking interneurons.
- ASR does not need predefined time points for removing spike-related-transient, but determines the start and end point of removal in each frequency band depending on changes in the power spectral density relative to the spike trough. This *temporal flexibility* ensures its usability for neurons with different shapes of action potentials and for variable action potentials of the same neuron.
- ASR is able to remove spike-related-transients that are not locked to the sharp rise of the spike and show variable timing. The variable timing of slow components is considered within up to one cycle for each frequency band. Adaptive timing of low frequency spike-related-transients makes the method suitable to not only detect high frequency oscillations but to provide a better signal to noise ratio for detecting low frequency oscillations.
- ASR is based on RMS matching, adaptive removal which makes it more robust to noisy signals than other threshold-based approaches.

We have implemented the ASR algorithm in Matlab and provide the commented code and a test dataset as free open source software on the website at www.accl.psy.vanderbilt.edu/resources/code/. The current implementation was made computationally efficient. Applying ASR to typical empirical datasets is achievable with standard computational resources. With the current implementation the adaptive artefact removal of 100 spikes takes on average 0.201 second for 100 spikes on a single core 2.8 GHz Intel Core i7 processor. The time varies depending on the variability of the spikes. The computational time requirements scale linearly with the number of spikes. There is no minimum number of spikes required for ASR and the number of spikes do not affect the ASR quality.

In summary, the described adaptive spike-artifact removal method provides an objective and principled method to estimate and remove confounding spike potentials across the major frequency bands of wideband local field potential data. We believe that this approach can proof to be essential in uncovering local circuit cell-to-circuit interactions that governs the local transformation and routing of information in brain circuits (Fries, 2015; Voloh and Womelsdorf, 2017; Womelsdorf et al., 2014a; Womelsdorf et al., 2014b).

## Acknowledgements

This research was supported by a grant from the Canadian Institutes of Health Research (T.W.) CIHR Grant MOP_102482 and by the National Institute Of Biomedical Imaging And Bioengineering of the National Institutes of Health under Award Number R01EB028161 (T.W.). The content is solely the responsibility of the authors and does not necessarily represent the official views of the Canadian Institutes of Health Research or the National Institutes of Health. We thank Mariann Oemisch and Seyed Ali Hassani for recording and preparing of the data.

